# Insertion sequence elements associated with *Staphylococcus epidermidis* evolution in persistent orthopaedic device-related infections

**DOI:** 10.64898/2026.05.21.726754

**Authors:** James C. Littlefair, Carolin M. Kobras, Virginia Post, Ben Pascoe, David J. Baker, Christoph Erichsen, Mathew Stracy, Fintan Moriarty, Samuel K. Sheppard

## Abstract

**Background:** *Staphylococcus epidermidis* is a major cause of orthopaedic device-related infections (ODRIs), which are often challenging to treat due to their extensive antimicrobial resistance (AMR) and biofilm formation. It has been hypothesised that *S. epidermidis* may rapidly adapt to the medical device niche, enhancing persistence, but direct evidence of within-host pathoadaptive evolution remains limited.

**Results:** To investigate within-host evolution during chronic infection by *S. epidermidis*, we analysed isolates from patients with confirmed ODRIs and used a rat infection model to examine the evolution of strains from two distinct epidemic lineages (ST2 and ST23). Our analysis revealed that the replicative transposition of insertion sequence (IS) elements within the accessory genome was the predominant mechanism of genetic diversification. This was largely driven by the IS256 family, which accounted for approximately 25% of all mutational events. However, other than SCC*mec* deletions resulting in the loss of *mecA,* no mutations, including those which exhibited parallel evolution, were predicted or observed to influence AMR or biofilm formation. These findings suggest that the strains investigated in this study, which already exhibited high-level multidrug resistance and biofilm-forming ability, were likely pre-adapted epidemic *S. epidermidis* clones well suited to establishing persistent ODRIs.

**Conclusions:** Our findings highlight the prominent role of IS elements in driving genetic diversification in *S. epidermidis*, underscoring the need for closer examination of their contribution to pathoadaptation during persistent infection.

## Background

*Staphylococcus epidermidis* is a proficient coloniser of human skin, where carriage is believed to be ubiquitous (1). Although it is not usually pathogenic in this context (2,3), the species has become increasingly associated with opportunistic life-threatening infections (4–6). This has been driven largely by its ability to colonise indwelling biomedical devices, such as those used in arthroplasties and trauma surgeries, which total at least 35–40 million procedures worldwide each year (7–11). Despite rigorous sterilisation protocols, the postoperative rate of orthopaedic device-related infections (ODRIs), periprosthetic joint infections, and fracture-related infections remains around 2% (12–14). However, incidence rates may exceed 50% in certain high-risk cases, such as open fracture surgeries (15). Approximately 20% of ODRIs and 30% of periprosthetic joint infections are attributed to *S. epidermidis*, often requiring both prolonged antibiotic therapy and invasive revision surgery to treat (7,9,16).

The pathogenesis of *S. epidermidis* is complex, governed by the dynamic interplay between the bacterium and the host (9,17,18). Several studies have shown that biofilm formation is a key driver of pathogenicity, with strong correlations with treatment failure (10,19,20). This is due to biofilms enhancing adhesion to indwelling devices, facilitating immune evasion, and reducing the efficacy of specific antimicrobial agents (10,21,22). Furthermore, antimicrobial resistance (AMR) is common in *S. epidermidis* (5,23–26). Resistance to β-lactams, such as oxacillin, has become so widespread – often exceeding 90% in certain clinical settings – that vancomycin has become recommended as a first-line agent for empiric treatment (7,26,27). Furthermore, studies conducted in Sweden, Poland, and Portugal have demonstrated that over 80% of clinical *S. epidermidis* strains exhibit multidrug resistance (MDR), defined as resistance to at least three classes of antibiotics (28–30).

While evidence remains mixed on whether significant genetic differences exist between clinical and carriage *S. epidermidis* isolates (29,31–35), the consistent overrepresentation of certain multilocus sequence types (STs), including ST2, ST5, and ST23, in opportunistic, persistent infections is strongly suggestive of pathoadaptation (24,29,36–39). It has been hypothesised that, in opportunistic pathogens such as *S. epidermidis*, pathoadaptation may occur both in the native commensal niche (pre-adaptation) as well as the secondary infection niche, with horizontal gene transfer (HGT) and mobile genetic elements playing a central role in this process (40). For example, in staphylococci, AMR determinants are frequently carried on plasmids (41) or chromosomally integrated mobile genetic elements such as the staphylococcal cassette chromosome *mec* (SCC*mec*) (42). Furthermore, there is increasing evidence that bacteriophages and transposons, including insertion sequence (IS) elements, contribute to pathoadaptation in staphylococci by mediating genome plasticity and HGT (43–47).

Compared to *Staphylococcus aureus*, very few studies have so far investigated the within-host evolution of *S. epidermidis* (33,48). To address this gap, we investigated *S. epidermidis* isolates collected longitudinally from persistent ODRIs, combining a novel experimental design with a bespoke variant-calling pipeline capable of detecting structural variants often overlooked by conventional single nucleotide polymorphism (SNP) and gene-centric methods of genomic surveillance. Specifically, two *S. epidermidis* strains isolated from patients with persistent ODRIs were inoculated into rats for 8 weeks without antibiotic treatment. This revealed that IS element insertions played a major role in within-host diversification, while large SCC*mec* deletions were associated with restored susceptibility to β-lactams. Overall, this study reveals a potentially underestimated role of IS elements in facilitating pathoadaptation in *S. epidermidis*, suggesting that greater focus on these elements could enhance our understanding of the evolutionary processes driving bacterial persistence in ODRIs.

## Methods

### Patient isolate collection

The collection of patient isolates was approved by the Ethikkommission der Bayerischen Landesärztekammer (accession 12063), and all patients provided written consent prior to participation. Detailed inclusion and exclusion criteria for the original cohort are available under accession NCT02640937 on clinicaltrials.gov (49) and in the Materials and Methods section of Morgenstern et al., 2016 (20) and Post et al., 2017 (50). Briefly, patients were prospectively enrolled if they were being treated at BGU Murnau, Germany, between November 2011 and September 2013 for a confirmed *S. epidermidis* infection following fracture fixation or prosthetic joint surgery. To minimise the risk of sampling contaminants, all bacterial samples were obtained from deep bone biopsies taken during the initial surgery after enrolment and subsequently from follow-up biopsies collected at the site of infection. Patient isolates have previously been analysed in Harris et al., 2020 (51). For the present study, longitudinal samples from ten patients were selected from the original cohort to generate the patient isolate collection (Table S1).

### Rat isolate collection

Ethical approval to use Wistar rats in this study was granted by the Graubünden Animal Commission under accession 32_2016. All procedures were performed in accordance with Swiss animal protection laws in an AAALAC-accredited facility. To produce this collection, two *S. epidermidis* isolates (SE13 and SE42), representative of isolates collected from patients 13 and 42, were each inoculated into 5 female Wistar rats using a tibial screw infection model previously described by Stadelmann et al., 2015 (52). The rats did not receive antibiotics at any time. After 8 weeks of infection, rats were euthanised and tibias were collected and homogenised. Homogenate was plated on Columbia agar with 5% sheep blood and incubated for 24-48 hours at 37°C. Up to ten colonies were harvested per rat, focussing on single colony separation (Table S1). All colonies were stored in 20% glycerol stocks at −80°C and revived as needed by streaking onto Columbia agar supplemented with 5% sheep blood, followed by incubation at 37°C for 24-48 hours at 37°C.

### Short read sequencing

All isolates were sequenced using Illumina^®^ short-read technology (Table S1). For DNA extraction, single colonies were picked from each plate, resuspended in tryptic soy broth (TSB), and grown in a shaking incubator overnight at 37°C at 180 rpm. 1.5 mL of overnight culture was subsequently centrifuged at 15,000 *g* for 2 minutes. The bacterial cell pellet was resuspended in 390 µL of Tris-EDTA (TE) buffer, to which 10 µL of lysostaphin in 20 mM, pH 5.2 sodium acetate buffer (5 mg mL^−1^) was added. This solution was then incubated for 2 hours at 37°C and 180 rpm. Following incubation, 10 µL of RNase A (10 mg mL^−1^) was added to the solution and left for 10 minutes. DNA was purified using the Maxwell^®^ RSC Cultured Cells DNA Kit, following the Maxwell^®^ RSC Automated DNA Purification protocol with Maxwell^®^ RSC software v3.0. DNA was quantified with a Promega Quantus™ Fluorometer using 5 µL of purified DNA according to the manufacturer’s instructions. DNA extractions were repeated until a minimum concentration of 40 ng µL^−1^ was achieved. DNA concentrations above 50 ng µL^−1^ were diluted with Milli-Q^®^ ultrapure water so that all final DNA concentrations were between 40 and 50 ng µL^−1^. 10 ng of DNA was used as input for library preparation. Libraries were prepared using the Illumina^®^ DNA Prep, (M) Tagmentation, kit with Unique Dual Index adapters, generating an insert size of approximately 350 bp. Pooled libraries were loaded at 750 pM onto an Illumina^®^ NextSeq™ 2000 using XLEAP-SBS™ P2 reagents for a paired-end 300-cycle run (2 × 150 bp).

### Long read sequencing

The two *S. epidermidis* strains used as the rat inoculum, SE13 and SE42, were long-read sequenced using Oxford Nanopore Technology™ (ONT). DNA extraction for long read sequencing used single colonies, which were resuspended in 200 µL of phosphate-buffered saline (PBS). 100 µL of this solution was then added to 20 mL of TSB in a sterile flask, which was incubated at 37°C and 180 rpm. Regular optical density at 600 nm (OD_600_) readings were taken by diluting 100 µL of liquid culture with 900 µL of TSB until an OD_600_ of 0.1 was achieved. 10 mL of liquid culture was then centrifuged at 3000 *g* for 10 minutes. The supernatant was discarded, and the bacterial cell pellet was resuspended in 0.5 mL of Zymo Research™ DNA/RNA Shield. DNA libraries were prepared with the ONT SQK-RBK114.96 kit using 200-400 ng of high molecular weight DNA. Pooled libraries were then loaded onto an ONT GridION™ using the FLO-MIN114 (R10.4.1) Flow Cell.

### Read processing and assembly

Using fastp v0.22.0 (53), Illumina^®^ short reads were deduplicated, adapter trimmed, quality trimmed (minimum average Phred quality of 30 in 4 bp sliding window in the 3’ direction), and filtered (minimum trimmed length of 71 bp and average Phred quality of 30). ONT long reads were length and quality-filtered using NanoFilt v2.8.0 (54) to a minimum length of 1000 bp and average Phred quality of 20. Processed short reads were assembled with SKESA v2.5.1 (55) using a k-mer length of 31. The processed short and long reads for the two strains inoculated into rats (SE13 and SE42) were used to generate hybrid assemblies using Trycycler v0.5.5 (56). For each strain, Trycycler *subsample* was used to create 12 read subsets at an approximate depth of 50×, based on an estimated genome size of 2.7 Mbp and 2.4 Mbp for strains SE13 and SE42, respectively. Four random subsets were then assembled using each of Flye v2.9.5-b180 (57), Miniasm and Minipolish v0.1.3 (58), and Raven v1.8.3 (59). Assemblies were clustered and processed with Trycycler *cluster*, *reconcile*, *multiple sequence alignment* (*msa*), and *partition* to generate consensus sequences, which were concatenated into a combined assembly. The combined assembly was polished using pypolca v0.3.1 (60,61) with processed short reads, and recircularised so that chromosomes began with the start codon of *dnaA*, while plasmids were orientated to start with the start codon of their respective resistance genes (*catA*, *qacA*, *qacB*, or *ermC*).

### Genomic characterisation

Species determination and multilocus sequence typing (MLST) of all isolates was performed with mlst v2.23.0 (62,63) using SKESA assemblies (Table S1).

Hybrid assemblies were annotated using the NCBI’s Prokaryotic Genome Annotation Pipeline (PGAP) v2024-07-18 (64,65) within a Docker v27.4.0 (66) environment. In addition, a curated set of protein sequences associated with AMR and biofilm formation (Table S4) were queried against the hybrid assemblies using tBLASTn v2.16.0 (67–69) with the following non-default parameters: *db_gencode* was set to 11 (the bacterial translation table), *comp_based_stats* was set to 0, and both *soft_masking* and *seg* were set to false. High-scoring pairs (HSPs) were sorted according to bit score and overlapping HSPs greater than 50 bp were filtered out unless they were assigned to the same gene query.

Hybrid assemblies for strains SE13 and SE42 were combined with 249 unique ST assemblies (Table S10) from the AllTheBacteria (70) database (aggregated releases 0.2 and 2024-08) to define core genome coordinates (alignment blocks shared between 100% of input strains) using CoreDetector v1.0.0 (71,72), applying a 5% divergence threshold and a minimum alignment length of 200 bp. Bacteriophage coordinates were identified (Table S13) using PHASTEST (73–75) in deep annotation mode. SCC*mec* typing was performed using SCC*mec*Finder (76) with default settings and putative SCC*mec* attachment (att) sites were identified using 13-mer sequences derived from the 3’ end of *orfX* (ribosomal RNA large subunit methyltransferase H, *rlmH*) within strains SE13 and SE42, excluding the base immediately upstream of the stop codon (Table S12). Due to the short and variable nature of SCC*mec* att sites, all possible combinations of variable positions between these two 13-mers were generated to create a set of query sequences. These queries were then searched against the hybrid assemblies for strains SE13 and SE42 using BLASTn with a word size of 4 bp and an E-value cutoff of 100. HSPs were filtered to retain only those ≥11 bp in length that were located between the start codons of *orfX* and *orfY* (putative tRNA-dihydrouridine synthase, *dus*).

### Variant-calling pipeline

To call variants, processed short reads were first mapped to their respective hybrid reference assemblies (inoculation strains SE13 or SE42) using bwa-mem2 v0.7.18-r1243-dirty (77,78) with a non-default mismatch penalty of 2. The resulting alignments were sorted and indexed with samtools v1.21 (79,80) to generate BAM files. These BAM files were used as input for InSurVeyor v1.1.2 (81) and Manta v1.6.0 (82), and were converted back to SAM format for input into QuickVariants v1.2.3 (83). InSurVeyor was run with default parameters. Manta was run using *configManta.py* followed by *runWorkflow.py* with default settings, and the resulting *diploidSV.gz* output was processed using *convertInversion.py*. QuickVariants was run with non-default parameters: *distinguish-query-ends* was set to 0, and *snp-threshold* and *indel-threshold* were configured to require a minimum read depth of 5 and a minimum allele frequency of 0.7. Variant calls from these pipelines were combined and characterised with a bespoke post-processing script, which is available on GitHub (Sheppard-Lab/RefToVariants). As it was noted that inoculation strains had diverged slightly from the isolates collected from patients 13 and 42, variants shared by all patient isolates but not strains SE13 and SE42 were excluded from subsequent analysis. Full lists of variants, including those that were excluded, can be found in Tables S2 and S3.

### Variant distribution analysis

Variants detected in patients and rats for both SE13 and SE42-related isolates were independently classified into mutually exclusive classes (IS element insertions, SNPs, InDels ≤7 bp, InDels >7 bp, SNP clusters, SCC*mec* deletions, and inversions) by the post-processing script, and additionally for genic variants, sub-classes (synonymous SNPs, missense SNPs, nonsense SNPs, in-frame InDels, out-of-frame InDels, and IS element insertions). Analysis of variant enrichment in the core and accessory genome was then performed at three levels: (1) total variant counts, (2) per variant class, and (3) per variant sub-class within genic regions. At each level, two-sided binomial tests (*binom.test*, base R v4.3.1) were used to evaluate whether the observed number of variants in the core genome deviated from the expected number under a random distribution. For levels 1 and 2, the expected proportion was based on the relative sizes of the core and accessory genome (Table S11) as determined by CoreDetector (71,72). For level 3, the expected proportion was based on the relative lengths of core and accessory genic regions, determined from the intersection between PGAP-annotated genes and CoreDetector-defined regions (Table S11). Bonferroni corrections (*p.adjust*, base R v4.3.1) were applied within levels 2 and 3 independently, with SE13 and SE42-related isolates treated as separate analyses throughout.

#### IS element family activity analysis

For each IS element insertion, IS element families were assigned automatically by the post-processing script using BLASTn v2.16.0 (67–69) with the inserted sequence as the query and the reference genome (SE13 or SE42) as the subject, identifying overlapping annotated features from PGAP-generated GFF files. These assignments were then manually curated to ensure each insertion was grouped within the correct IS element family. To assess the relative transposition activity of IS element families, Fisher’s exact tests (*fisher.test*, base R v4.3.1) were performed for each family independently. For each family, insertion counts and intact copy numbers were compared against the pooled counts of all other families combined, forming a 2×2 contingency table. Families with no intact copies in the original genome were excluded from the analysis. Tests were performed separately for SE13 and SE42-related isolates, and Bonferroni corrections (*p.adjust*, base R v4.3.1) were applied within each strain independently to account for multiple comparisons.

### AMR disk diffusion testing

Resistance to oxacillin, gentamicin, and erythromycin was assessed using disk diffusion tests for isolates with variants in associated resistance determinants (Table S8). Overnight cultures were grown in triplicate and diluted in TSB to a final OD_600_ of 0.1. Diluted cultures were spread on tryptic soy agar (TSA), and 6 mm Oxoid™ antimicrobial susceptibility disks containing defined amounts of antibiotic (oxacillin, 1 µg per disk; gentamicin, 10 µg per disk; erythromycin, 15 µg per disk) were applied with adequate spacing. Standard European Committee on Antimicrobial Susceptibility Testing (EUCAST) clinical breakpoints (84) were used to interpret inhibition zone diameter (oxacillin, 20 mm; gentamicin, 22 mm; erythromycin, 21 mm) and classify isolates as susceptible or resistant.

### Biofilm assay

The biofilm assay performed in this study was adapted from Stepanovic et al., 2007 (85), Morgenstern et al., 2016 (20), and Post et al., 2025 (86). Briefly, for each isolate, overnight cultures were diluted to an OD_600_ of 0.1 in TSB, and again diluted 1:100 in TSB supplemented with 1% glucose. This solution was then inoculated on a 96-well culture plate, with three technical replicates per overnight culture, and statically incubated for 20 hours at 37°C. Culture OD_600_ readings were taken using a VANTAstar^®^ microtiter plate reader before discarding the contents, rinsing with PBS in three cycles, and drying at 55°C for 60 minutes. Gram’s crystal violet (0.1% w/v in distilled water) was then added for 15 minutes, and washed three times with deionised water until the run-off water was free of stain. After drying for 30 minutes at room temperature and resolubilising the cells with 95% ethanol for 30 minutes at room temperature, final biofilm OD_600_ readings were obtained using a VANTAstar^®^ microtiter plate reader (Table S9).

Technical replicate biofilm OD_600_ readings that deviated by more than two standard deviations from the sample mean were suspected of contamination and excluded for subsequent analyses (Table S9). Biofilm-forming ability for each isolate was classified using an optical density cutoff (OD_c_), defined as the mean OD_600_ of the negative control plus three standard deviations. The four levels of the scale were defined as follows: no biofilm (OD_600_ ≤ OD_c_), weak biofilm (OD_c_ < OD_600_ ≤ 2×OD_c_), moderate biofilm (2×OD_c_ < OD_600_ ≤ 4×OD_c_), and strong biofilm (OD_600_ > 4×OD_c_). In addition, biofilm formation was analysed using a linear mixed-effects (LME) model in R (*lme4* v1.1.37). The model included negative control-adjusted biofilm OD_600_ as the response variable, with isolate and negative control-adjusted culture OD_600_ as fixed effects, and 96-well plate and biological replicates as random effects. Technical replicates were nested within overnight culture replicates in the LME model to account for the experimental design. The model was then used to perform *post hoc* pairwise comparisons based on estimated marginal means (EMMs) in R (emmeans v1.11.2) with a Bonferroni correction (*p.adjust*, base R v4.3.1).

## Results

### Detection of genetically diverse isolates in patient ODRIs provides evidence of polyclonal infection or clonal replacement

To investigate the diversity of *S. epidermidis* strains involved in patient ODRIs, we performed whole-genome sequencing and multilocus sequence typing (MLST) on isolates collected longitudinally from 10 patients with either osteomyelitis or infected endoprostheses (Fig. 1a). In this collection, the most frequently observed sequence type (ST) was ST2, detected in five patients, followed by ST5 and ST23, each detected in three patients (Fig. 1b). These findings are consistent with previous reports describing ST2, ST5, and ST23 as globally disseminated, nosocomial lineages (24,29,36–39). With the exception of patient 9, who harboured only clade B isolates (ST328 and ST595), all patients carried isolates belonging exclusively to clade A (Fig. 1c), the predominant clade found on skin as well as clinical isolates (3,31,50).

**Fig. 1:**
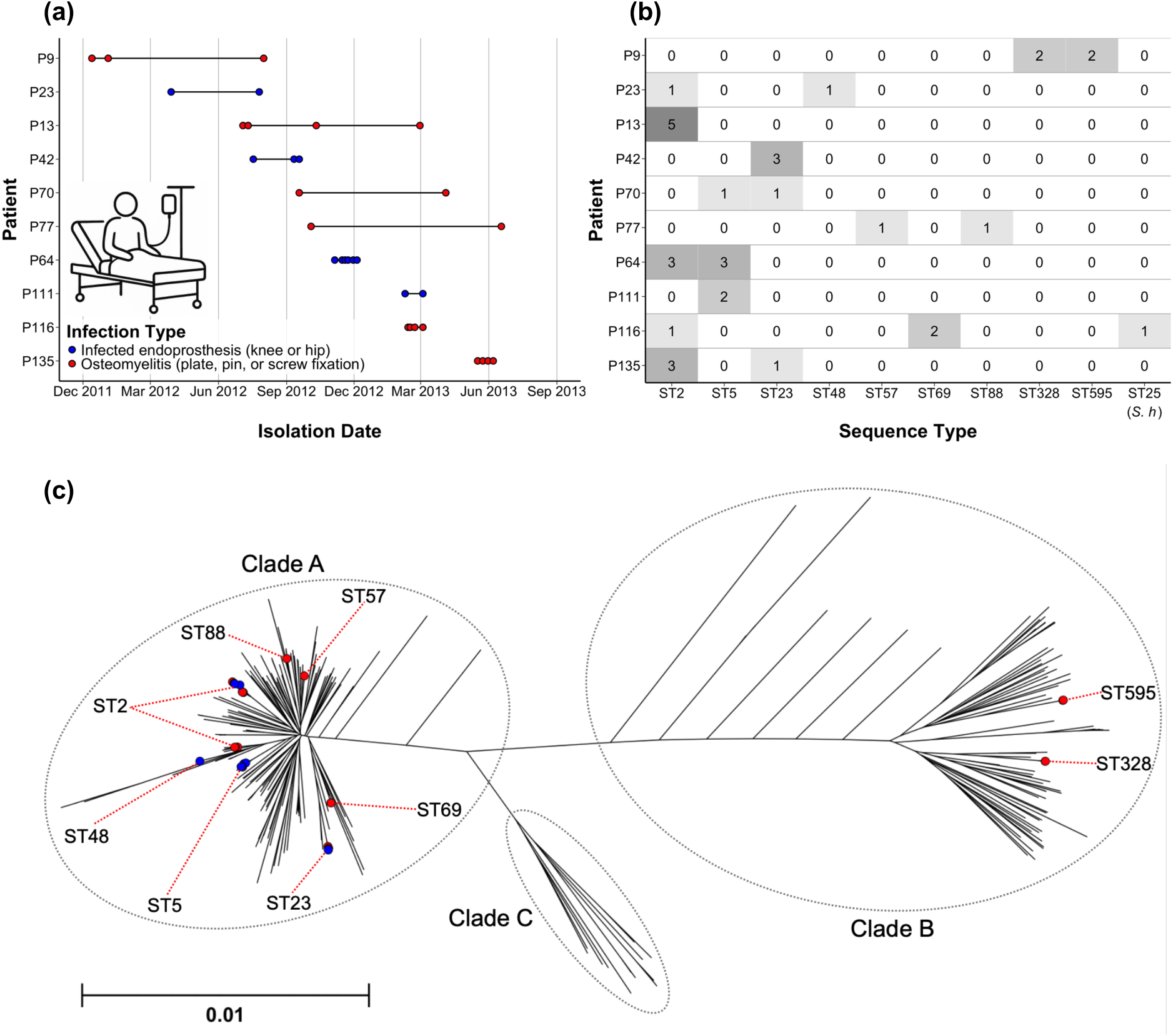
*Staphylococcus epidermidis* exhibits substantial genetic diversity within and between patient ODRIs. **(a)** Longitudinal sampling of ten patients with ODRIs between December 2011 and June 2013 at a specialist medical facility in Murnau, Germany. Each point represents the date on which a patient isolate was collected from deep bone biopsy sampling. **(b)** Multilocus sequence types (STs) of isolates collected from each patient. Counts denote the number of patient isolates belonging to each ST. All isolate STs were assigned using the *S. epidermidis* schema, except for ST25, which was a *S. haemolyticus* isolate (indicated as *S. h*). **(c)** Population structure of *S. epidermidis* including patient isolates. K-mer-derived core genome distances were determined between a set of 283 diverse isolates, including 249 population samples (Table S10), each representing a unique ST, and 34 patient samples (Table S1). Distances represent the proportion of non-identical core genome k-mers as computed with pp-sketchlib. Patient isolates are indicated with circular tips: red for osteomyelitis and blue for infected endoprostheses.

Despite the substantial clade A bias, considerable diversity was observed at the ST level, with only three patients colonised by the same ST across all timepoints (Fig. 1b). For the remaining seven patients, two distinct STs were observed, with one patient also yielding a strain of *Staphylococcus haemolyticus*, suggesting a polymicrobial infection involving at least two coagulase-negative staphylococci. These findings highlight the potential contribution of both polyclonal and polymicrobial infection to *S. epidermidis*-associated ODRIs. However, in the absence of parallel sampling, the detection of multiple strains could alternatively reflect repeated reintroduction of skin flora following disruption of the epidermal barrier, for example during revision surgery or through fistula formation. As our primary aim was to investigate within-host evolution, we selected isolates from two patients with monoclonal infections for further study, where confidence was high that these strains truly represented persistent infection. These isolates were obtained from patients 13 and 42, and belonged to ST2 and ST23, respectively (referred to herein as SE13 and SE42).

### Within-host evolution revealed a mutational landscape dominated by IS element insertions and SNPs in patients and rats

To determine if mutations observed during patient ODRIs were characteristic of such infections, we inoculated each of two *S. epidermidis* strains associated with monoclonal infections into five rats and harvested colonies after eight weeks (Fig. 2a). These strains, designated SE13 and SE42, were representative of isolates collected from patients 13 and 42, which allowed for direct comparison between variants observed in patient and rat isolates (Tables S2 and S3).

**Fig. 2:**
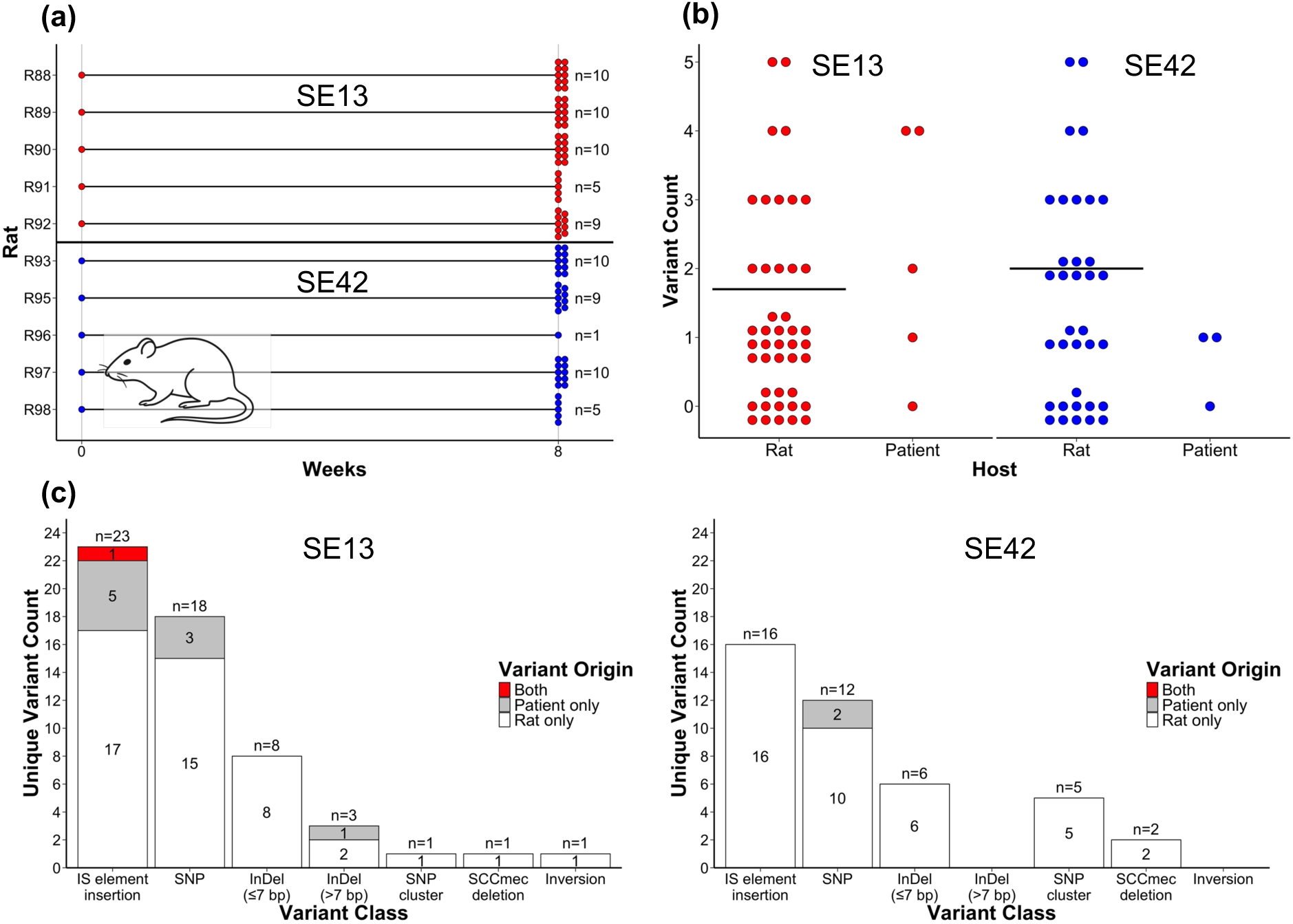
Within-host diversification of SE13 and SE42-related isolates was predominantly driven by IS element insertions and SNPs. **(a)** Inoculation and sampling of strains SE13 and SE42 in rat ODRI model. SE13 and SE42 were each inoculated into five rats and multiple colonies were harvested after 8 weeks of infection. The number of *S. epidermidis* isolates collected from each rat which were closely genetically related (<10 variants) to the inoculation strains are indicated. **(b)** Rat and patient isolate counts by the number of variants they contained relative to strains SE13 and SE42. The mean number of variants detected in rat-evolved isolates are indicated with horizontal black bars **(c)** Unique variant counts by variant class. Variants were considered to be unique if they occurred at different loci or involved different variant classes at the same locus. Stacked bars indicate the number of unique variants detected exclusively in rats (white), exclusively in patients (grey), or in both hosts (red).

After eight weeks of infection, rat isolates accumulated between 0 and 5 variants, with a mean of 1.7 and 2 variants for SE13 and SE42-derived isolates, respectively (Fig. 2b). Although all patient isolates exhibited variant counts in this range, mutation rates could not be accurately inferred because the strains sampled at the earliest patient timepoints were not common ancestors of subsequently sampled isolates, suggesting diversification had occurred prior to initial sampling. In both strain backgrounds, IS element insertions were the most frequently observed variant class, followed closely by single nucleotide polymorphisms (SNPs), with other classes of mutations being relatively infrequent (Fig. 2c). Although direct comparison of variant classes between patients and rats was limited by the small sample size of patient isolates, the variant classes observed in patients were among the most frequently detected in rats (Fig. 2c).

### Mutations including IS element insertions were enriched in the accessory genome

Pinpointing where mutations occur within genomes can provide key insight into the evolutionary processes underlying within-host evolution. To investigate this, we combined variants from both patient and rat isolates and examined their distribution across the core and accessory genome of SE13 and SE42 (Fig. 3a). The core genome, generated from the shared sequence between 251 genetically diverse *S. epidermidis* isolates (Table S10), including strains SE13 and SE42, was estimated to span approximately 1.7 Mbp and be 91% genic – protein-coding genes and ncRNAs (Table S11), as annotated with PGAP (65). For SE13 and SE42, core regions represented 62% and 65% of the total genome length, respectively (Table S11). Despite this, only 19 of 55 (35%) variants detected for SE13 and 18 of 41 (44%) variants detected for SE42 were located in core regions (Fig. 3a), meaning, in both strain backgrounds, variants were roughly three times more abundant in the accessory genome than in the core – significantly fewer than expected at random based on the relative sizes of the core and accessory genomes (two-sided binomial tests, SE13: *p* = 4.1×10^−5^, SE42: *p* = 0.0079).

**Fig. 3:**
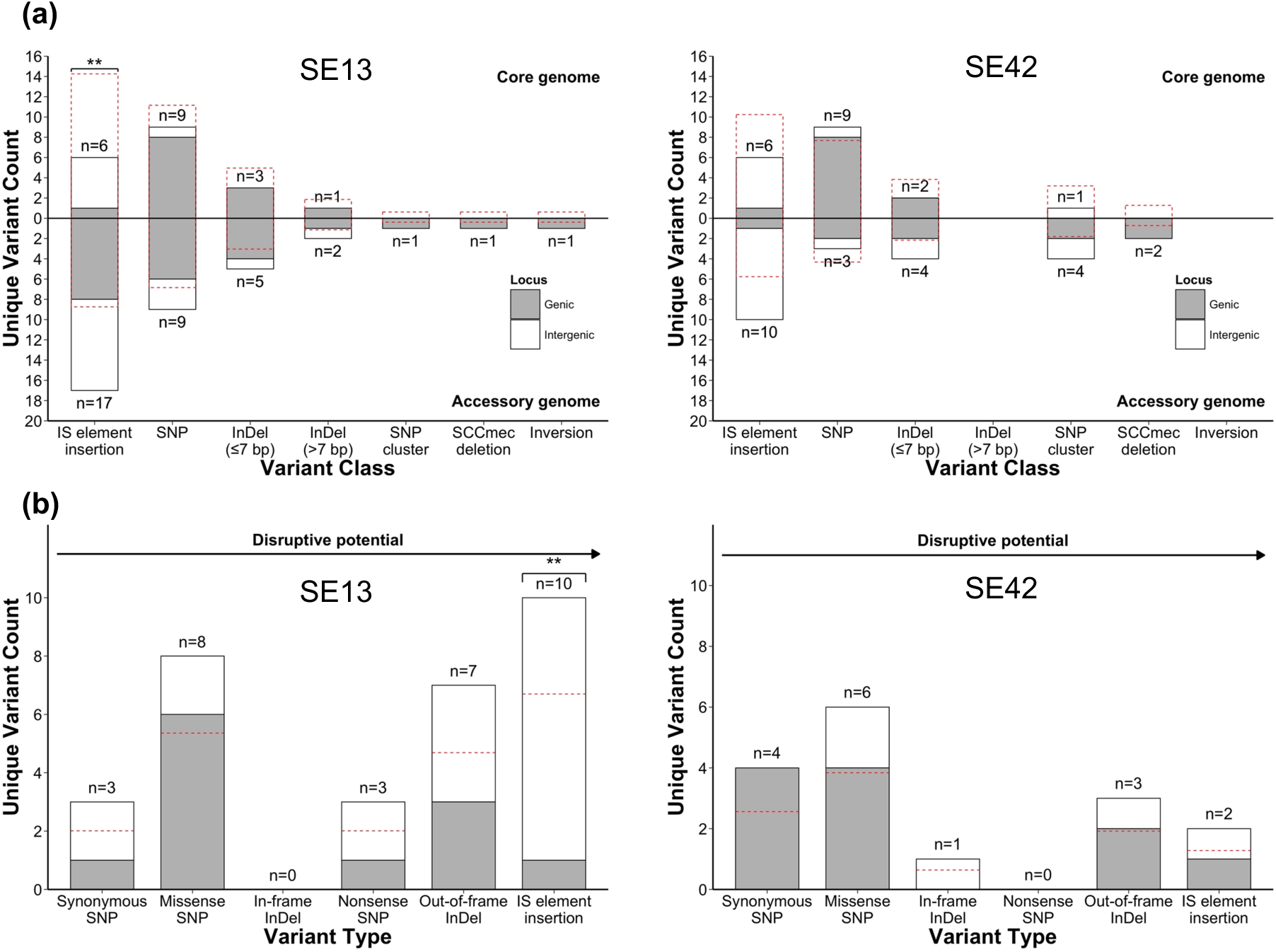
Genomic distribution of variants reveals overall bias for accessory regions, particularly for IS element insertions. **(a)** Unique variant counts by variant class in the core and accessory genome. Bars above the x-axis represent unique variants in the core genome, while bars below represent unique variants in the accessory genome, with total counts indicated. Stacked bar segments distinguish variants in protein-coding genes (grey) from those in intergenic regions (white). Red dashed bars indicate the expected distribution of unique variants at random based on the relative lengths of the core and accessory genome. Statistically significant deviations from this distribution are indicated with asterisks (* *p*_adj_ < 0.05, ** *p*_adj_ < 0.01, *** *p*_adj_ < 0.001). **(b)** Unique genic (protein-coding) variant counts by genic variant subclass (variant type). Stacked bars represent the total number of unique genic variants detected, with grey corresponding to core genic regions and white to accessory genic regions. Red dashed lines indicate the expected distribution of variants under a random distribution based on the relative total lengths of core and accessory genic regions. Grey bars below the dashed line indicate enrichment of variants in accessory genic regions. Statistically significant deviations from the expected number of variants are indicated with asterisks, using the same *p*_adj_ thresholds described above.

The uneven distribution of variants in the core and accessory genome was further examined by variant class, revealing that most were overrepresented in the accessory genome (Fig. 3a). However, in part due to the small number of variants in most classes, only IS element insertions observed in SE13 reached statistical significance (two-sided binomial test, *n* (core) = 6, *n* (accessory) = 17, expected proportion (core) = 0.62, *p*_adj_ = 0.0054). Compared to other variant classes, SNPs were more evenly distributed between core and accessory regions and were also more frequently observed in genic, protein-coding regions (Fig. 3a). The relative paucity of other variant classes within these core genic regions likely reflects strong purifying selection, as disruption of these genes would often be deleterious because they are typically associated with essential housekeeping functions (87–89). Consistent with this hypothesis, genic variant sub-classes with greater disruptive potential, such as IS element insertions, out-of-frame InDels, and nonsense SNPs, were generally underrepresented in core genic regions compared to those with lower disruptive potential – such as synonymous SNPs, missense SNPs, and in-frame InDels (Fig. 3b). Similarly to their general overrepresentation in the accessory genome, IS element insertions in SE13 were also significantly overrepresented in accessory genic regions (two-sided binomial test, core *n* = 1, accessory *n* = 9, expected core proportion = 0.67, *p*_adj_ = 0.0016).

### IS256 demonstrated elevated activity consistent with replicative transposition

Given that IS element insertions were the most commonly observed variant class, we next assessed the relative activity of different IS element families. This revealed IS256 as the most active family in both SE13 and SE42, accounting for over 60% of new IS element insertions despite comprising only approximately 20% of intact IS elements in the original genomes (Table 1), representing significantly elevated transposition activity relative to its genomic abundance in both strains (Fisher’s exact test, SE13: OR = 6.9, *p*_adj_ = 0.0048; SE42: OR = 7.4, *p*_adj_ = 0.039). In contrast, some IS element families showed low insertion frequencies relative to their abundance. For instance, IS3 had 8 intact copies in the original genome of strain SE13 and IS110 had 12 intact copies in the original genome of strain SE42, but each was observed only once in new insertions. Consistent with previous studies which have reported extremely low excision rates for IS elements such as IS256 (90,91), no IS element deletions were detected for any family, strongly suggesting they predominantly employ replicative mechanisms of transposition.

**Table 1:** IS element insertion frequencies for SE13 and SE42.

| Strain | Family | Original Copies |  | New Insertions |  |
| --- | --- | --- | --- | --- | --- |
|  |  | Intact (n) | Defective (n) | Frequency (n) | Frequency (%) |
| SE13 | IS3 | 8 | 1 | 1 | 4.3 |
|  | IS110 | 13 | 0 | 7 | 30.4 |
|  | IS200 | 2 | 1 | 0 | 0 |
|  | IS256 | 7 | 0 | 15 | 65.2 |
|  | IS257 | 4 | 1 | 0 | 0 |
|  | IS1182 | 0 | 5 | 0 | 0 |
|  | Total | 34 | 8 | 23 | 100 |
| SE42 | IS3 | 1 | 0 | 0 | 0 |
|  | IS110 | 12 | 0 | 1 | 6.3 |
|  | IS200 | 2 | 2 | 0 | 0 |
|  | IS256 | 4 | 0 | 10 | 62.5 |
|  | IS257 | 3 | 0 | 0 | 0 |
|  | IS1182 | 1 | 5 | 5 | 31.3 |
|  | Total | 23 | 7 | 16 | 100 |

### IS element insertions exhibited parallel evolution consistent with adaptation

The unique experimental design of this study, incorporating multiple rat replicates, enabled the identification of variants arising independently through parallel evolution. Although no identical variants were detected among biological replicates for SE42, three IS element insertions, two involving IS110 and one involving IS256, occurred at identical or closely adjacent genomic loci in SE13-evolved isolates, indicating parallel evolution had occurred.

The IS element insertion showing the strongest evidence for parallel evolution was an IS110 insertion observed in isolates collected from three of five rats and the original patient. This insertion occurred within an unnamed AAA family ATPase gene located on a large 147 kbp prophage with over 99% sequence identity to ΦSepi-HH1 (33). Notably, insertion bias of IS elements toward prophage regions has been previously reported in *Escherichia coli*, where such events are thought to contribute to prophage immobilisation through gene inactivation (92).

A second IS110 insertion exhibiting parallel evolution was observed in isolates collected from two of five rats. This insertion occurred approximately 50 bp downstream of the *clpB* gene, within what is likely its 3’ untranslated region (UTR), based on transcriptome analyses in *S. aureus* (93). Although the function of the *clpB* 3’ UTR remains unclear, ClpB itself is known to play an important role in the bacterial stress response (94), suggesting that this insertion may represent an adaptive mechanism for modulating *clpB* expression under conditions of stress.

The final IS element insertion associated with a mutation hotspot involved IS256, detected in isolates from two rats within a gene encoding an unnamed hypothetical protein. Unlike the aforementioned IS110 insertions, which occurred at identical positions between biological replicates, this hotspot comprised of two insertions separated by 18 bp. This pattern may indicate a lower target-site specificity for IS256, compared to other IS families, potentially contributing to its elevated transpositional activity. Consistent with this observation, IS256 is known to target AT-rich regions but does not have a well-conserved consensus target sequence (95).

### Mutations did not affect biofilm formation for SE13 or SE42

Given that previous studies have reported within-host evolution in staphylococci leading to increased biofilm formation (48,96), we investigated the effect of variants carried by rat-evolved isolates on biofilm production. Although no variants were detected within or nearby to known biofilm-associated genes, we tested 14 isolates carrying novel candidate mutations (10 evolved from SE13 and 4 evolved from SE42) using a biofilm assay (Table S9). This assay revealed no significant differences in biofilm formation between evolved isolates and their respective original strains (Fig. 4). However, it did reveal that strain SE42 was a significantly stronger biofilm former *in vitro* than SE13, producing approximately 5-fold more biofilm after 20 hours of growth (LME model with EMMs, estimate = 0.57; 95% CI = 0.52-0.61; *p*_adj_ < 0.0001). Analysis of 36 biofilm-associated genes (Table S4) revealed little to no variation in gene presence, sequence, or intactness between the two strains. For instance, both strains carried the *icaADBC* operon, which is strongly associated with biofilm formation. The most notable differences were that SE42 possessed the *bhp* (Bap Homologue Protein) and *sdrG* (Serine-aspartate Repeat-containing Protein G) genes, which were absent from SE13, suggesting that these genes may contribute to the enhanced biofilm formation observed for SE42.

**Fig. 4:**
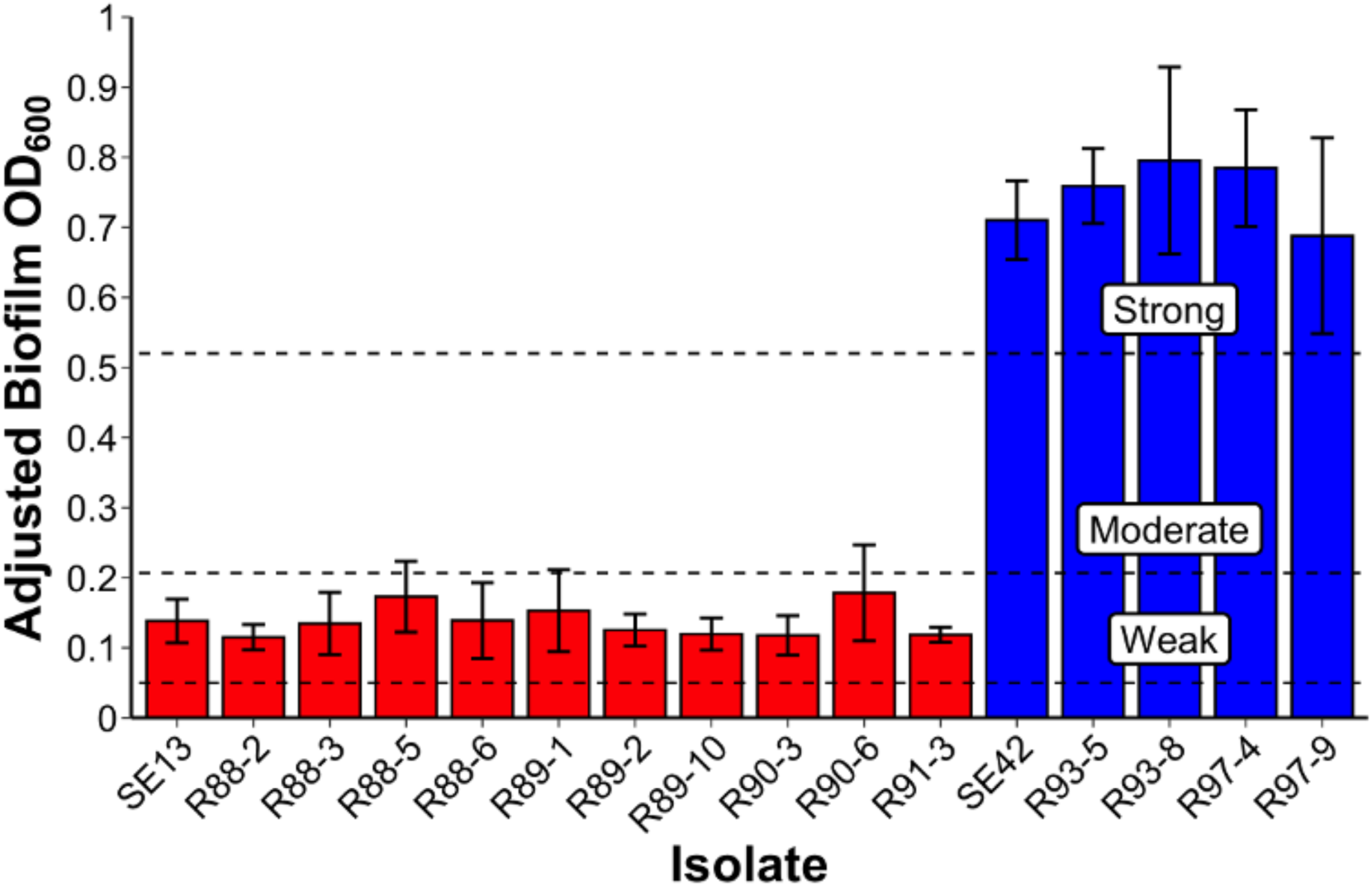
Biofilm assay revealed SE42 was a significantly stronger biofilm former than SE13, but a shift toward increased biofilm formation in rat-evolved isolates was not observed. Negative control-adjusted biofilm OD_600_ by sample. OD_600_ measurements were normalised by subtracting the mean OD_600_ of negative control wells (containing no sample). Bar heights represent the mean adjusted OD_600_, with error bars indicating ±1 standard deviation. Red bars correspond to SE13-evolved isolates, while blue bars correspond to SE42-evolved isolates. Horizontal dashed lines indicate the minimum adjusted OD_600_ thresholds used to classify isolates as weak, moderate, and strong biofilm formers, as described in detail in the Materials and methods section.

### SCC*mec* deletions restored susceptibility to oxacillin in SE13 and SE42

AMR plays a critical role in the pathogenesis of *S. epidermidis*, with many resistance determinants located on mobile genetic elements. Among these, SCC*mec* is perhaps the most well-known. This element typically carries *mecA* or *mecC*, which encode alternative penicillin-binding proteins that confer resistance to β-lactam antibiotics such as oxacillin (97).

In this study, three large deletions were observed in rat-evolved isolates within SCC*mec* – two in strain SE13 and one in strain SE42 (Fig. 5a). Two of these SCC*mec* deletions resulted in the loss of *mecA*, *aadD1* (associated with aminoglycoside resistance), and *bleO* (associated with bleomycin resistance). Disk diffusion tests confirmed that the loss of *mecA* restored susceptibility to oxacillin but the loss of *aadD1* did not diminish resistance to gentamicin, likely due to the presence of the *aac(6’)-Ie-aph(2”)-Ia* gene in both genomes (Fig. 5b; Table 2). Bleomycin susceptibility was not assessed. All three deletions occurred between short repeat sequences with high homology to the canonical attB attachment site of SCC*mec* located within *orfX*. This suggests an important role for these repeats in facilitating recombination, which is mediated by Ccr recombinases encoded within SCC*mec* itself. The abundance of such repeats throughout the element likely contributes to its high propensity for recombination, explaining why large deletions were observed only within SCC*mec* and not elsewhere in the genome.

**Fig. 5:**
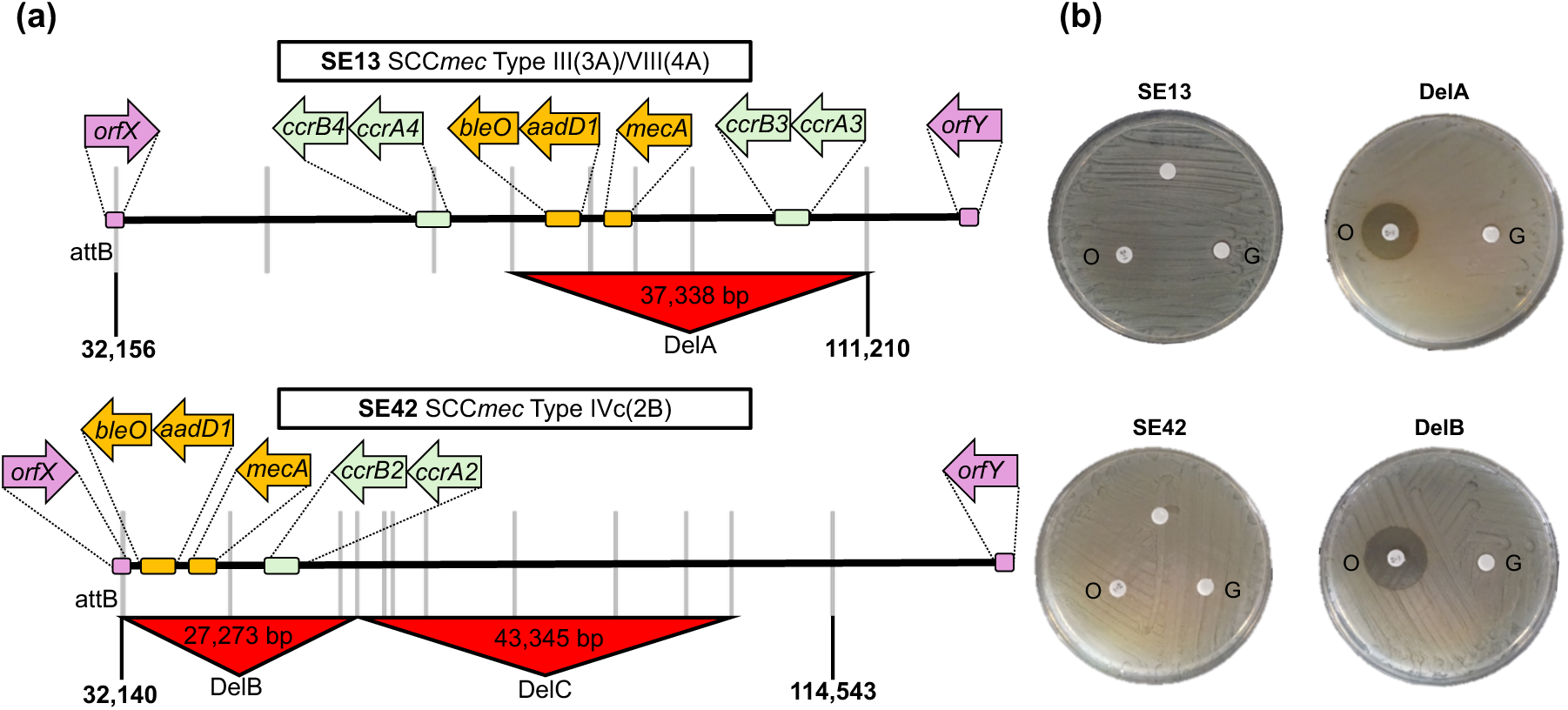
SCC*mec* deletions in SE13 and SE42-evolved isolates restored sensitivity to the β-lactam antibiotic oxacillin. **(a)** SCC*mec* structure in SE13 and SE42. SCC*mec* spans are shown as chromosomal coordinates between the attB attachment site located in *orfX* (*rlmH*) and the attachment site located immediately downstream of *orfY* (*dus*). The positions of all attachment sites with homology to attB are shown as vertical grey bars. SCC*mec* deletions, which all occurred between these attachment sites, are shown as red triangles, with the size of the deletion indicated. The relative positions of antimicrobial resistance genes and CCr recombinases are shown in orange and green, respectively, with the direction of the arrow indicating the orientation of these genes with respect to *orfX*. All genes were predicted to be full length functional copies except for *addD1* in SE13 which existed as a truncated fragment at approximately 61% coverage. Genes shown in expanded arrows are not to scale. **(b)** Disk diffusion tests confirming restored sensitivity to β-lactams (oxacillin) but not aminoglycosides (gentamicin). Disks containing oxacillin and gentamicin are indicated with capital letters (O and G, respectively).

**Table 2:** AMR genes detected in strains SE13 and SE42.

| Strain/s | Gene | Antibiotic | Location |
| --- | --- | --- | --- |
| SE13, SE42 | *mecA* | Oxacillin | Chromosome |
|  | *gyrA*<sup>84F</sup> | Ciprofloxacin | Chromosome |
|  | *aac(6')-Ie-aph(2'')-Ia* | Gentamicin | Chromosome |
|  | *rpoB*<sup>471E,527M</sup> | Rifampicin | Chromosome |
|  | *ermC* | Erythromycin | Plasmid |
| SE13 | *dfrA*<sup>99Y</sup> | Trimethoprim | Chromosome |
| SE42 | *catA* | Chloramphenicol | Plasmid |

As rats were not administered antibiotics throughout infection, the observed loss of resistance in some evolved isolates may not be unexpected if they confer substantial fitness costs in this context. However, it was notable that all mutations involving known resistance genes occurred within the SCC*mec* element – the three deletions described above, as well as an independent IS110 insertion within *bleO* in strain SE13, likely rendering it non-functional. Although both strains were highly multidrug resistant (MDR), being resistant to at least six commonly used antimicrobial agents (Table 2), variants were not detected in any other resistance gene. This suggests that most resistance determinants in these strains remained genetically stable throughout the 8-week infection period, irrespective of whether they were located chromosomally or on plasmids, even in the absence of selective antibiotic pressure.

## Discussion

*S. epidermidis* remains a major public health concern due to its prominence in ODRIs, the incidence of which continues to rise year on year (98–100). Given the substantial health and economic burden associated with these infections (8,101), numerous studies have sought to elucidate the factors that enable this species to transition from a ubiquitous skin commensal to a successful opportunistic pathogen – for example, by comparing infection isolates with those carried asymptomatically (29,31,35,102–105). However, to date, few studies have examined the within-host evolutionary processes that drive pathoadaptation during persistent infection (33,106). In this study, we addressed this gap by characterising samples collected longitudinally from ten patients with ODRIs and conducting a parallel evolution experiment in rats.

Our analysis of longitudinally collected patient isolates revealed that most patients (7 of 10) harboured infection isolates with diverse STs, which primarily belonged to the clade A phylogroup. We also observed the presence of a *Staphylococcus haemolyticus* isolate in one patient, indicative of polymicrobial infection. Although data on the frequency of polyclonal *S. epidermidis* infections remains limited, they have previously been observed in intravascular catheter infections and ODRIs (51,107). In contrast, polymicrobial infections involving *S. epidermidis* have been reported extensively (108–112). Interestingly the skin microbiome has been shown to harbour highly diverse *S. epidermidis* lineages, primarily belonging to clade A (3,31). We therefore hypothesise that ‘normally sterile sites’ associated with ODRIs such as bones and joints may commonly become infected with multiple founder lineages from the skin microbiome during surgery or subsequently through reintroduction of skin flora following disruption of the epidermal barrier, resulting in polyclonal and polymicrobial infections.

To the best of our knowledge, this study is the first to comprehensively quantify the frequencies of different variant types arising during within-host evolution of *S. epidermidis* during persistent ODRIs. Our analyses revealed that replicative IS element insertions were the primary drivers of diversification in this context, accounting for approximately 40% of all mutational events in strains evolved from SE13 and SE42 across patients and rats. Notably, the IS256 family alone was responsible for about 25% of all events, demonstrating markedly elevated transpositional activity in comparison to other IS element families. IS256 has previously been identified as a key molecular marker of invasive *S. epidermidis* lineages, comparable to markers associated with AMR and biofilm formation (33,105,113–115), and interestingly has also been shown to drive within-host evolution in *S. aureus* bacteraemia (116), where up to three new IS256 insertions per isolate were observed, even in the absence of other mutations. These findings suggest that, contrary to the outdated view of IS elements as genomic parasites, IS elements may play an important role in the pathoadaptation of this species, as has been previously demonstrated in *Enterococcus faecium* and *Enterococcus faecalis* (117,118).

The pathoadaptive potential of IS elements has been previously explored in *S. epidermidis and S. aureus*, with particular focus on biofilm-associated phenotypic variation (90,119). In *S. epidermidis*, the *icaADBC* operon, which is strongly associated with biofilm formation, was initially identified as a phase-variable insertion hotspot for IS256 (90). However, subsequent studies challenged this view, reporting that IS256 insertions within the *icaADBC* operon were rare (120,121) and that, when they were present, the rate of precise excision was orders of magnitude lower than that of insertion (91). Consistent with these later findings, we did not detect any new IS element insertions within or immediately adjacent to genes associated with biofilm formation, including within the *icaADBC* operon. Although this could simply reflect selective pressure to maintain biofilm-forming ability during persistent infection (122), the complete absence of excision events elsewhere in the genome strongly suggests that IS elements in *S. epidermidis* primarily mobilise through replicative transposition, which does not strongly support a role in phase variation.

Compared with the highly diverse skin microbiome where *S. epidermidis* natively resides (3,123,124), “normally sterile sites” associated with ODRIs (such as joints and bones) are characterised by low microbial abundance and limited taxonomic diversity (125–127). In this isolated niche, opportunities for HGT are presumed to be extremely limited (128). Therefore, where opportunities for HGT are more limited, we hypothesise that IS elements may play an especially important role in generating genetic novelty through rapid accessory gene inactivation, a mechanism previously shown to drive evolution in the microbiota of the human gut (129), and evidenced in this study by the three IS element insertions exhibiting parallel evolution. These insertions occurred within protein-coding regions and 3’ untranslated regions of genes, suggesting that gene suppression – via open reading frame disruption or mRNA destabilisation – may be a key mechanism by which IS elements facilitate pathoadaptation in this hostile niche. Supporting this hypothesis, an IS110 insertion was observed within the AMR gene *bleO*. Since the rats were not exposed to antibiotics, it is plausible that *bleO* became redundant during the evolution of this isolate, and that its inactivation conferred a selective advantage if its expression imposed an unnecessary metabolic cost.

In addition to IS element insertions, we also detected three large independent SCC*mec* deletions in rat-evolved isolates, a phenomenon which has been previously observed in patient-matched *S. epidermidis* isolates (33,130). Two of these deletions involved AMR determinants, including *mecA*, which restored oxacillin sensitivity in these evolved isolates. Interestingly, no other large deletions were found elsewhere in the genome of any evolved isolate, and other AMR determinants carried by strains SE13 and SE42 were not involved in mutation, supporting previous findings that identified SCC*mec* as a hotspot for sequence instability (130–132). We demonstrated that in both strains these deletions occurred between short repeat sequences with high identity to canonical attB insertion site of SCC*mec*, suggesting that these were probably mediated by the cassette-specific Ccr recombinases encoded within the element itself.

The emergence and global dissemination of epidemic *S. epidermidis* clones are of particular concern due to their ability to establish persistent ODRIs. Our findings indicate that the two strains analysed in detail in this study using the rat infection model, SE13 and SE42, likely represent pre-adapted epidemic *S. epidermidis* lineages capable of establishing such persistent ODRIs. Both SE13 and SE42 belonged to ST2 and ST23, respectively, lineages previously identified as globally disseminated, hospital-acquired clones (24,36–39). Consistent with this, SE13 and SE42 exhibited high-level MDR, were *icaADBC*-positive biofilm formers, and carried highly active IS256 elements. These traits, which are hallmarks of successful nosocomial *S. epidermidis* clones (102), likely contribute to their persistence and adaptability within hospital environments and during chronic infection. As discussed above, opportunities for HGT are more likely to be limited at the site of infection than on the skin. Consequently, as we did not observe gene acquisition through HGT in evolved isolates, it is plausible that many of these genetic traits were acquired within the native skin microbiota during the evolution of these epidemic clones, indirectly conferring selective advantages under the hostile conditions encountered during infection.

Although great care was taken to ensure the accuracy of the results presented in this study, some limitations in the study design should be acknowledged. Firstly, both strains analysed exhibited all the hallmarks of pre-adapted epidemic lineages. While this may be advantageous for studying the evolution of pre-adapted healthcare-associated lineages, it does not provide insight into how strains may initially undergo pathoadaptation, for example, within the skin microbiome or site of infection. Additionally, the limited level of genome annotation available for many loci, particularly in non-coding regions, made it challenging to infer the potential phenotypic effects of many IS element insertions. Finally, the sampling strategy – especially for patient isolates – introduced some constraints. The small number of patient isolates, together with the collection of only a single isolate per timepoint, made it difficult to assess within-host strain heterogeneity in patients and to draw broad comparisons with the evolutionary patterns observed in rats.

## Conclusions

Overall, our results reveal that IS elements, particularly IS256, are predominant drivers of within-host diversification of *S. epidermidis* during persistent ODRIs. IS element insertions as well as SCC*mec* element deletions likely facilitate gene inactivation, which may allow infecting strains to rapidly adapt to the hostile isolated niche of “normally sterile sites”. Although we did not identify shifts toward increased biofilm formation in evolved isolates, we identified pre-existing MDR and biofilm-forming ability in the strains analysed, which belonged to known nosocomial lineages of *S. epidermidis*, consistent with pre-adaptation. Considering this, future efforts should be directed towards understanding the within-host evolution in both native sites of colonisation such as the skin as well as within the site of infection, with particular focus on mobile genetic elements including IS elements as drivers of such adaptation.

## Declarations

### Ethics approval and consent to participate

The collection of patient isolates was approved by the Ethikkommission der Bayerischen Landesärztekammer (accession 12063), and all patients provided written consent prior to participation. Ethical approval to use Wistar rats in this study was granted by the Graubünden Animal Commission under accession 32_2016.

### Availability of data and materials

All data, software, and code required to replicate this study are referenced within the article or provided in the Supporting information. Raw sequence data are deposited in the National Center for Biotechnology Information (NCBI) Sequence Read Archive (SRA) under BioProject accession PRJNA1354684.

### Authors’ contributions

James C. Littlefair, Conceptualisation, Data curation, Formal analysis, Investigation, Methodology, Project administration, Software, Visualisation, Writing – original draft | Carolin M. Kobras, Conceptualisation, Data curation, Formal analysis, Investigation, Methodology, Resources, Writing – review and editing | Virginia Post, Conceptualisation, Data curation, Investigation, Methodology, Resources, Writing – review and editing | Ben Pascoe, Data curation, Formal analysis, Methodology, Writing – review and editing | David J. Baker, Investigation | Christoph Erichsen, Investigation, Resources | Mathew Stracy, Funding acquisition, Resources, Writing – review and editing | Fintan Moriarty, Conceptualisation, Data curation, Investigation, Methodology, Resources, Writing – review and editing | Samuel K. Sheppard, Conceptualisation, Funding acquisition, Methodology, Project administration, Resources, Supervision, Writing.

## Acknowledgements

Not applicable.

